# EGFR-driven profibrotic signals in AKI-to-CKD transition are blocked by TNF inhibition: Opportunities for etanercept treatment

**DOI:** 10.1101/2021.04.29.442002

**Authors:** Mai M Abdelmageed, Eirini Kefaloyianni, Akshayakeerthi Arthanarisami, Fatima Zohra Khamissi, Jeffrey J. Atkinson, Andreas Herrlich

## Abstract

Proximal-tubule-cell EGFR activation mediates tubule cell proliferation and repair early after kidney injury, while sustained EGFR activation causes kidney fibrosis. Inflammation is a key driver of AKI-to-CKD transition and fibrosis, but mechanisms of EGFR-driven profibrotic responses are not well understood. In a mouse model of AKI-to-CKD transition and CKD progression, we show that EGFR-inhibition significantly reduced kidney expression of many immunoregulatory molecules already by day two after injury, including the potent inflammatory cytokine tumor-necrosis-factor (TNF). Single nuclei RNA-sequencing analysis showed that macrophages were among the main early cellular sources of TNF in the injured kidney. *In vitro*, EGFR activation in macrophages increased macrophage TNF expression, while EGFR inhibition *in vivo* reduced kidney macrophage accumulation, as early as two days after injury. Thus, profibrotic EGFR signaling increases kidney TNF levels both directly and indirectly. TNF inhibition did not alter tubule EGFR activation and in contrast to EGFR inhibition did not reduce early macrophage accumulation in the kidney, suggesting that TNF does not promote early infiltration of immune cells in the kidney, but rather regulates profibrotic functions of kidney and/or immune cells. TNF inhibition with etanercept *in vivo* in AKI-injured mice downregulated a number of cytokines, including TNF itself. This cytokine downregulation overlapped with EGFR inhibition, and additional non-overlapping downregulated cytokines shared the same functions, predicting that TNF inhibition would prevent AKI-to-CKD transition like EGFR inhibition. Indeed, TNF-inhibition with etanercept reduced AKI-induced fibrosis to the same degree as the EGFR-inhibition, while the combination of both treatments showed no additive effect. In conclusion, our results identify TNF as a downstream effector of profibrotic EGFR activation and motivate the examination of TNF pathway inhibition in human AKI or CKD.

**Translational statement:** Proximal tubule epidermal-growth-factor (EGFR) activation in mice and likely also in humans drives inflammation in the kidney after acute-kidney-injury (AKI) and causes chronic-kidney-disease (CKD) with fibrosis in a process termed AKI-to-CKD transition. Recent retrospective data shows that patients treated with TNF inhibitors show decreased incidence and progression of CKD. Our current work shows that TNF inhibition in mice is equally as effective as EGFR inhibition in preventing AKI-to-CKD transition and fibrosis. Thus, our results have translational potential and may stimulate examination of short-term TNF inhibition in AKI to prevent AKI-to-CKD transition or possibly of longer-term TNF inhibition in CKD to prevent CKD progression.

## Introduction

Inflammation is a major driver of fibrosis and progression of chronic-kidney-disease (CKD) resulting in progressive loss of kidney function. Similarly, acute kidney injury (AKI) can lead to persistent pro-fibrotic inflammation (AKI-to-CKD transition), in particular when the insult is severe, repetitive or chronic ^1,2^. In AKI and CKD macrophages represent important drivers of profibrotic inflammation ^3^. We previously showed that injured proximal-tubule-cells (PTC) in mice and humans upregulate a-disintegrin-and metalloprotease-17 (ADAM17)-dependent release of the profibrotic epidermal-growth-factor-receptor (EGFR) ligand amphiregulin (AREG) ^4^. ADAM17- or AREG-PTC-knockout (KO) strongly reduced PTC-EGFR activation, kidney macrophage accumulation, inflammation and fibrosis after AKI in mice ^4,5^, identifying sustained ligand-induced PTC-EGFR activation as a key proinflammatory and profibrotic event in the development of fibrosis after AKI (AKI-to-CKD transition). The protective effect of EGFR-inhibition or of PTC-EGFR-KO in injury-induced fibrosis was first reported by the Friedlander laboratory and these findings were confirmed and extended by others ^6–10^. However, the mechanisms by which EGFR drives inflammation in AKI-to-CKD transition and fibrosis remain ill-defined.

In humans, CKD progression is strongly associated with increased serum levels of tumor-necrosis-factor (TNF), a potent type 1 inflammatory cytokine, and its soluble receptors TNFR1 and TNFR2 ^11–13^, which may act as decoy receptors for TNF. A recent large retrospective propensity-matched cohort study ^14^ and a smaller study ^15^ suggested that anti-TNF biologics, such as etanercept (TNFR2-Fc), reduce incidence and progression of CKD in patients with rheumatoid arthritis. In rodents only limited data is available regarding the role of TNF-inhibition in kidney injury and profibrotic inflammation. Early kidney injury markers decreased when rats subjected to ischemia-reperfusion-injury (IRI) were pretreated with TNF-inhibition by etanercept, but no late fibrosis outcomes were measured ^16^. TNF-inhibition with soluble pegylated TNFR1 in rats reduced early fibrosis markers after UUO ^17^, but another UUO study in TNF-global-KO mice showed increased fibrosis^14^. The dominant cellular sources of TNF in the injured kidney were not determined in these studies. Several studies that reported effects of the human TNF-inhibitor infliximab in rodent kidney injury models focusing on early injury ^18–21^ have been called into question since it was shown that infliximab does not bind to rodent TNF ^22^, suggesting that observed effects were independent of TNF neutralization. Taken together, these findings warrant reevaluation of the role of TNF-inhibition in kidney injury and fibrosis.

We investigated immuno-regulatory effects of EGFR inhibition *in vivo* during the early critical phase of AKI-to-CKD transition when profibrotic inflammation is first established. Using single nucleus RNAseq data, we identify TNF released by macrophages as an early critical pro-fibrotic event downstream of EGFR. We show that EGFR-inhibition and TNF-inhibition significantly overlap in downregulating kidney cytokines after AKI, in particular of TNF, and have the same anti-fibrotic effects. Mechanistically, EGFR inhibition downregulates kidney TNF after AKI by blocking the recruitment of TNF-producing macrophages, whereas TNF inhibition neither affects EGFR activation, nor does it prevent macrophage recruitment, but achieves its anti-fibrotic effects by blunting the action of soluble TNF released by immune cells recruited by EGFR signaling. These findings, together with the retrospective observation that TNF-inhibition reduces incidence and progression of CKD in some patients, predict a possible role to TNF inhibition in preventing AKI-to-CKD transition and the progression of fibrosis in humans that could be prospectively tested in the clinic.

## Methods

### Animal studies

For all studies adult (8-12 week-old) male mice were used in accordance with the animal care and use protocol approved by the Institutional Animal Care and Use Committee of Washington University School of Medicine, in adherence to standards set in the Guide for the Care and Use of Laboratory Animals (8th edition, The National Academies Press, revised 2011). Ischemia for 21 min at 37°C was induced in both kidneys using the flank approach as previously reported ^49^. Sham operations were performed with exposure of both kidneys, but without induction of ischemia.

### Animal treatments

Erlotinib hydrochloride was purchased from LC Laboratories (Woburn, MA, USA). It was dissolved in a vehicle consisting of 0.5% methyl cellulose + 1% tween 80 in water at a concentration of 8 mg/ml and stored in aliquots at −20 □C. Erlotinib was administered to animals at a dose of 80mg/kg/day via gavage ^8^. Murine etanercept was kindly provided by Pfizer at a concentration of 12.5 mg/ml, it was diluted using PBS and injected to animals at a dose of 10 mg/kg i.p. twice per week. Mice were randomly assigned into groups and received treatments for 2 or 28 days (starting from the surgery day until day 1 or day 27 respectively) as follows; *Sham*: received both Erlotinib via gavage daily and Etanercept i.p. twice per week, *and* had a sham operation instead of bilateral ischemia operation, *Vehicle/Vehicle* (*V/V*): received the vehicle used for dissolving Erlotinib via gavage daily and was injected vehicle PBS i.p. twice per week, *Etanercept* (*ET*): received Etanercept i.p. twice per week and vehicle used for dissolving Erlotinib via gavage daily, *Erlotinib* (*ER*): received Erlotinib gavage daily and PBS i.p. twice per week, *Combination* (*COMB*): received both Erlotinib gavage daily and Etanercept i.p. twice per week. Mice were sacrificed on day 2 or day 28. A group of mice received fluorescently labeled etanercept (the VRDye™ 549 kit from LI-COR was used for labeling according to manufacturer’s instructions) to confirm delivery of the drug in the kidney interstitium.

### Renal function and histology

Serum creatinine was assessed by an LC-mass spectrometry-based assay at the O’Brien Core Center for Acute Kidney Injury Research (University of Alabama School of Medicine, Birmingham, Alabama, USA). BUN levels were measured using the DiUR100 kit (Thermo Scientific) according to the manufacturer’s instructions. GFR was measured by monitoring the clearance of FITC-sinistrin (Fresenius-Kabi, Linz, Austria) using the MediBeacon Transdermal GFR Monitor (MediBeacon GmbH, Mannheim, Germany). Briefly, the Transdermal GFR Monitor was attached to the shaved area on the mouse back for 2 min then FITCsinisterin was injected to mice at a dose of 75 mg/Kg intravenously via the retro-orbital sinus. The fluorometer was allowed to record the change in FITC-sinisterin fluorescence over 1 hr. The device was then removed and connected to PC to download the recorded measurements. GFR was calculated using one compartmental kinetics fitting allowing direct conversion from the FITC-sinisterin elimination half-life obtained from the exponential excretion phase of the curve using a conversion factor. GFR (μL/min/25gBW) = 3654.2/ t_1/2_ (FITC-sinistrin) (min).

Kidney histology was examined in formalin-fixed sections. For each staining, 10 images per kidney throughout the tissue were collected for blinded quantification. Fibrosis severity was quantified in kidney cortex by measuring the Picrosirius red-stained area using ImageJ software ^50^.

### Immunofluorescence staining

Immunofluorescence staining of the kidney was performed on frozen sections following standard protocols. In brief, cryo-sections of tissue in OCT were washed with PBST (0.05% (v/v) Tween-20 in PBS) for 5 min at room temperature and permeabilized with 1% SDS in PBS for 5 min, followed by three 5 min washes with PBST. After 30 min incubation in blocking buffer (5% BSA and 5% goat serum in PBS), sections were incubated with primary antibodies overnight at 4oC, followed by three 5 min washes with PBST, 1 h incubation with secondary antibodies (where appropriate), three 5 min washes with PBST and mounting in DAPI-containing medium (Vector Laboratories). The primary Abs included: αSMA (FITCconjugated, Sigma-Aldrich, #F3777), fibronectin (Abcam, #ab2413) and CD31 (BD Biosciences #550274). For each kidney staining, 10 images per kidney throughout the tissue were collected for blinded quantification. Images were analyzed using the ImageJ software.

### Kidney lysate preparation and western blot

Whole kidney lysates were prepared as previously described^4^ and analyzed by standard western blot techniques. The primary antibodies used were from Cell Signaling Technology (p-EGFR Y1068 #2234) or from Abcam (GAPDH #ab181602).

### Cytokine profile assay

The proteome profiler array was performed using the Mouse XL Cytokine Array Kit (#ARY028, R&D systems, Minneapolis, USA) following the manufacturer’s instructions. Briefly, kidneys were homogenized in PBS to which protease inhibitors and 1 % Triton-X were added. Assay membranes were incubated with blocking buffer for 1 hr at room temperature and then with the lysates overnight at 4 □C. After washing, membranes were incubated with the detection antibody cocktail for 1 hr at room temperature. After another wash, membranes were incubated with Streptavidin-HRP for 30 min at room temperature, washed again and exposed to the Chemi reagent mix. Images were captured using the ChemiDoc Imager (BioRad) and analyzed using the ImageLab software (BioRad). The pixel density of each spot was determined using the reference spots and signal density was compared between different samples to determine the relative changes in the analyte levels between the different samples. Only cytokines reduced or increased by at least 30% with respect to vehicle were considered as downregulated or upregulated respectively. GO enrichment analyses for Biological Processes and transcriptional regulatory networks (TRRUST) were performed using Metascape ^51^. Venn diagrams were generated using BioVenn ^52^. Heatmaps were produced using GraphPad Prism, version 9.0 (GraphPad Software Inc.).

### Bulk RNA sequencing

Total RNA was isolated from mouse kidneys using the TRIzol reagent (Invitrogen) following the manufacturers’ instructions. Total kidney RNA samples were sequenced by Novogene Corporation Inc and the results were analyzed using GraphPad Prism, version 9.0 (GraphPad Software Inc.). Single cell preparation for RNA sequencing and mass cytometry: Kidneys were minced into small pieces (<1mm^3^*) and* incubated in tissue dissociation buffer (1 mg/ml Liberase TM, 0.7 mg/ml Hyaluronidase, 80U/ml DNAse in PBS) for 30 min in 37°C. Single cells were released from the digested tissue by pipetting 10 times and the cell suspension was filtered through a 70 μm sieve (Falcon). 10% Fetal Bovine Serum (FBS) was added to stop the enzymatic reaction. Cells were collected by centrifugation (300 g at 4°C for 5 min) and resuspended in red blood cell lysis buffer (155mM NH4Cl, 10mM KHCO3, 1mM EDTA, pH 7.3) for 1 min at room temperature. After washing in PBS, cells were either used fresh for analysis by scRNAseq or were cryopreserved in liquid nitrogen vapor in freezing media (50% fetal bovine serum, 40% RPMI 1640, 10% DMSO) until analysis by mass cytometry.

### Single nucleus RNA sequencing analysis

A publicly available dataset was used for analysis ^26^. The online tool for single cell data analysis of the Humphreys laboratory (Kidney Interactive Transcriptomics http://humphreyslab.com/SingleCell/) was used to analyze the expression of TNF in different kidney cell types at day 2 post IRI.

### Mass cytometry

Single cell preparations were analyzed by mass cytometry as previously described ^53^. Briefly, cells were labeled using a previously validated and titrated antibody cocktail for surface markers ^53^ (all antibodies conjugated by the manufacturer-Fluidigm) diluted in Fluidigm MaxPar Cell Staining Buffer (CSB) (1 hour at 4 °C). After two washes in CSB, cells were fixed in 2% PFA for 20 min at room temperature, washed, stained with MaxPar Intercalator-IR (Fluidigm) and filtered into cell strainer cap tubes. Data was then acquired on a CyTOF2/Helios instrument (Fluidigm) and analyzed using the CytoBank software.

### Statistics

All results are reported as the mean ± SEM. Comparison of 2 groups was performed using an unpaired, 2-tailed t-test or a Pearson correlation analysis where appropriate. Comparison of 3 groups was performed via ANOVA and Tukey’s post hoc test. Statistical analyses were performed using GraphPad Prism, version 9.0 (GraphPad Software Inc.). A *P* value of less than 0.05 was considered significant.

## Results

### EGFR inhibition reduces kidney TNF and other inflammatory cytokines that regulate leukocyte migration, adhesion and activation in AKI-to-CKD transition

We used an AKI-to-CKD severe bilateral renal ischemia-reperfusion-injury (IRI) model for our studies. To conclusively establish that this model leads to significant progressive loss of glomerular-filtration-rate (GFR), we evaluated mice after bilateral IRI or sham operation for a period of 6 months (180 days) (**Figure 1A**). Bilateral IRI caused significant AKI with serum blood-urea-nitrogen (BUN) and creatinine elevations on Day 1 after AKI, that returned to baseline after 1 month (**Supplemental Figure 1**). GFR, assessed trans-dermally by detecting the excretion of injected FITC-sinistrin ^23^, also returned to normal by 1 month after injury (**Figure 1B**). This occurred due to renal compensatory mechanisms and despite the fact that severe fibrosis is observed at this time point in AKI-injured animals, as we and others previously documented ^4,5,8,10^, and we also demonstrate later in Figure 5. To assess whether established fibrosis at 1 month would lead to progressive renal failure, we measured GFR over 6 months. GFR declined progressively in AKI-injured animals starting 3-4 months after injury, leading to a total loss of 40% of baseline GFR by six months after AKI as compared to sham (**Figure 1B**). Serum BUN and creatinine measured over the same time period were normal (**Supplemental Figure 1**), highlighting known insensitivity of these markers for determination of early kidney failure. Thus, our rodent AKI model behaves as predicted from studies in patients with AKI that progress to CKD (AKI-to-CKD transition) ^1,2^.

**Figure 1:**
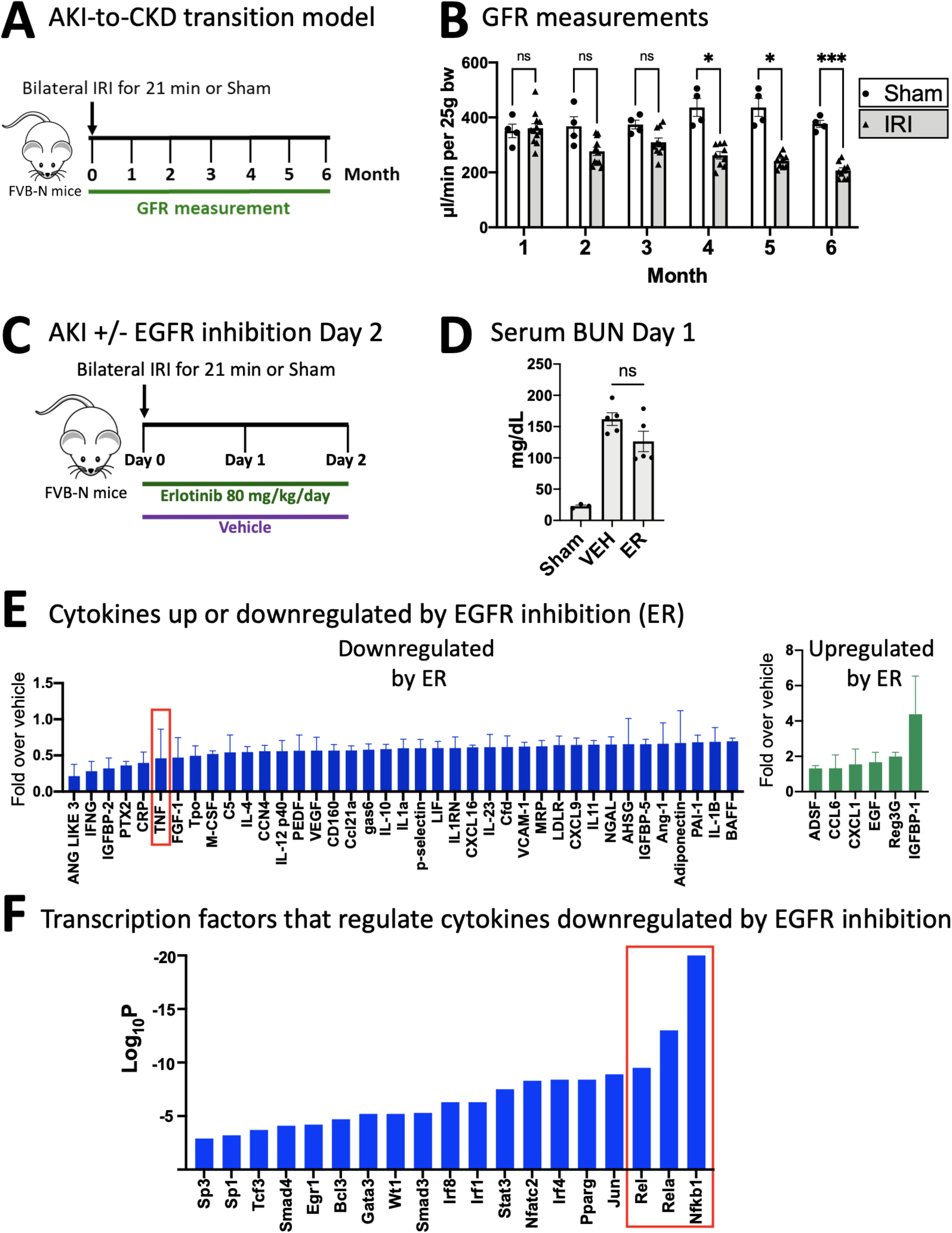
EGFR inhibition reduces kidney TNF and other inflammatory cytokines that regulate leukocyte migration, adhesion and activation in AKI-to-CKD transition. **A**. Experimental scheme of AKI-to-CKD model. **B**. Monthly glomerular filtration rate (GFR) measurements in Sham and AKI mice over 6 months. GFR decreases starting at 3-4 months after AKI. **C**. Experimental scheme of either vehicle or erlotinib treatment for 2 days after AKI. **D**. Serum BUN Day 1 (VEH: vehicle treatment; ER: erlotinib treatment). **E**. Cytokines downregulated (left blue column graph) or upregulated (right green column graph) by erlotinib (ER) treatment. TNF is highlighted in red rectangle. **F**. Enrichment analysis of transcription factors known to regulate cytokines downregulated by erlotinib treatment. NFκB family members are highlighted in red rectangle. *P < 0.05; ***P < 0.001; ns: non-significant

To gain a better understanding of EGFR’s immunomodulatory functions, we performed proteomic analysis of kidney tissue lysates for the expression of 98 proinflammatory/profibrotic cytokines, chemokines and related molecules (in the following simply referred to as cytokines). We compared IRI-injured mice treated with vehicle or the EGFR inhibitor erlotinib (ER) on Day 2 after injury (**Figure 1C**). On Day 2 after injury, phosphorylation of EGFR in the kidney, and in particular in tubule cells, is very significantly elevated ^4,8,24^. Erlotinib (ER) did not affect initial kidney injury, as shown by BUN elevations appropriate for the degree of injury and comparable to vehicle controls at day 1 (**Figure 1D**). However, EGFR inhibition very strongly reduced protein levels of a large number of cytokines. Top scoring downregulated cytokines are shown in **Figure 1E**. In particular, ER treatment strongly affected cytokines involved in regulation of leukocyte migration (GO term analysis, **Supplemental Table 1**). Only a small number of cytokines were increased by ER treatment (**Figure 1E**). The potent type I cytokine tumor-necrosis-factor (TNF), a major regulator as well as target of the NFκB family ^25^, was among the most strongly downregulated cytokines in ER treated mice (**Figure 1E**, highlighted by a red rectangle). Enrichment analysis of transcription factors known to regulate the expression of any of the cytokines downregulated by ER treatment indeed strongly downregulated transcription factors of the NFκB family (**Figure 1F**, highlighted by a red rectangle). A summary of our transcription factor enrichment analysis is available in **Supplemental Table 2**. We thus hypothesized that TNF is an important downstream target of EGFR signaling after kidney injury.

### EGFR increases kidney TNF levels by driving accumulation of macrophages in the injured kidney and enhancing macrophage TNF expression

To better understand the role of EGFR activation in kidney TNF expression, we first aimed to determine cellular sources of TNF in the injured kidney. Using a publicly available single nuclei RNA sequencing (snRNAseq) dataset ^26^, we found that macrophages represent a major source of kidney TNF early (day 2) after ischemic injury (**Figure 2A**). We tested the hypothesis that EGFR activation may affect kidney TNF levels either indirectly, by modulating the accumulation of TNF-expressing macrophages in the injured kidney, and/or by acting directly on macrophages to stimulate TNF expression. Most macrophages that accumulate in the kidney after injury are known to be derived from recruited circulating monocytes that express chemokine-receptor-2 (CCR2), the monocyte/macrophage receptor of the cytokine Ccl2 (also called monocyte-chemoattractant-protein-1 (MCP-1)), and blockade of CCR2 ameliorates kidney fibrosis ^27–29^. Bulk mRNA sequencing analysis showed that erlotinib (ER), as compared to vehicle control (VEH) significantly reduced kidney levels of the global immune cell marker CD45, the macrophage/dendritic cell marker F4/80, and dramatically reduced levels of CCR2; the macrophage marker CD64 also showed a trend to reduction (**Figure 2B**). Mass cytometric analysis confirmed reduced interstitial macrophage accumulation in the kidney already on day 2 after AKI (**Figure 2C**). Thus, EGFR inhibition strongly reduces TNF protein in the kidney by blocking ingress of inflammatory cells that represent a main source of TNF after injury. We next tested whether EGFR signaling might directly affect TNF production in macrophages, by stimulating RAW264.7 macrophages with EGFR ligand *in vitro* and examining TNF mRNA expression. We found that the low affinity EGFR ligand amphiregulin (AREG), which we previously showed is highly upregulated in the kidney after AKI and mediates profibrotic EGFR functions in AKI-to-CKD transition ^4,5^, induces transcriptional upregulation of TNF in macrophages, as determined by qPCR (**Figure 2D**). Taken together these results suggest that EGFR activation regulates TNF levels in the kidney, in two ways: indirectly by driving attraction and accumulation of macrophages in the kidney early after AKI, when they represent the main TNF expressing cell-type, and possibly also directly, by enhancing TNF expression levels in macrophages via stimulation of EGFR signaling. We thus hypothesized that TNF could mediate at least part of EGFR’s profibrotic functions.

**Figure 2:**
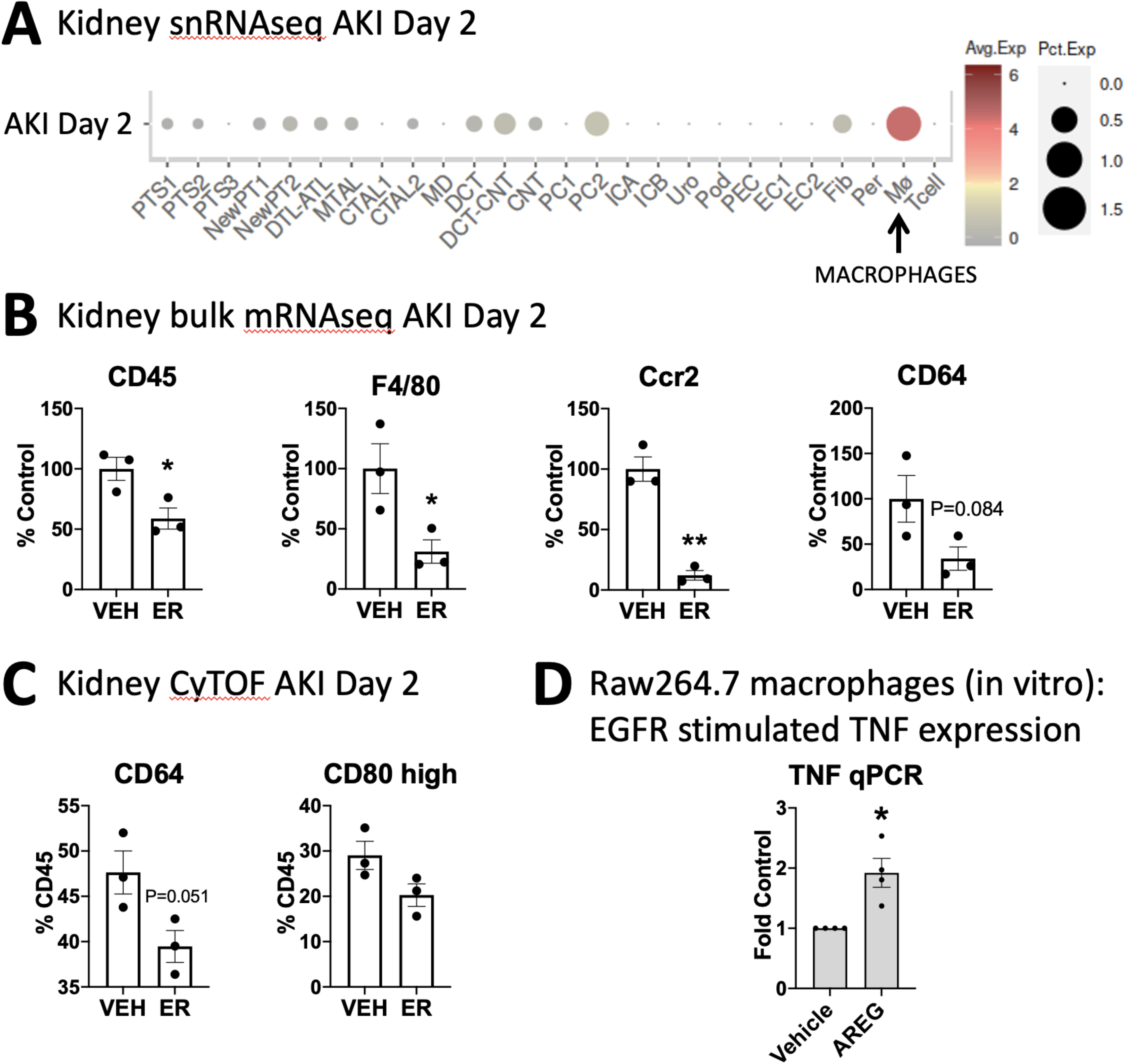
EGFR increases kidney TNF levels by driving accumulation of macrophages in the injured kidney and enhancing macrophage TNF expression. **A**. snRNAseq analysis of AKI-kidneys at Day 2 after injury shows macrophages as the main TNF expressing cells. **B**. Bulk mRNA sequencing analysis of kidney at Day 2 after AKI in vehicle (VEH) or erlotinib (ER) treated animals. Expression of the general immune cell marker CD45, of the macrophage/dendritic cell marker F4/80, and of CCR2, an important recruitment signal for macrophage-precursor monocytes, are all significantly reduced by ER, suggesting ER drives recruitment and accumulation of macrophages in the injured kidney (n=3); the macrophage marker CD64 also showed a trend to reduction. **C**. Mass cytometry analysis of kidney Day 2 after AKI. ER significantly reduces the number of CD64 positive macrophages after AKI. A similar trend is seen for the CD80 macrophage marker (n=3). **D**. qPCR analysis of Raw264.7 macrophages stimulated by vehicle or the EGFR ligand amphiregulin (AREG), showing increased TNF expression after EGFR stimulation (n=4). *P < 0.05

### TNF inhibition does not interfere with kidney EGFR activation or macrophage accumulation in the kidney early after injury

To determine whether TNF could indeed mechanistically act as a key driver of fibrotic inflammation downstream of EGFR, we tested the effect of TNF inhibition on kidney EGFR activation and on EGFR-driven profibrotic downstream events *in vivo*. Etanercept (ET) is a TNFR2-Fc conjugate that acts as a decoy receptor that sequesters soluble TNF. For our experiments, we used the murine version of the drug (murine Tnfr2-Fc, Pfizer ^30,31^) to ensure specificity and efficiency of mouse TNF protein binding. Etanercept (ET) was effectively delivered into the interstitial compartment of the kidney, as shown by injection of fluorescently-labeled ET (**Supplemental Figure 2A**). ET was found partially colocalized with vascular endothelial cell marker CD31 and partially isolated in the extravascular interstitial space between tubule cells (**Supplemental Figure 2B**). First, we examined EGFR phosphorylation at early time points in etanercept treated mice, using cortical kidney lysates, which are highly enriched for proximal tubule. TNF-inhibition (ET) did not significantly affect kidney tubule EGFR phosphorylation *in vivo*, whereas EGFR-inhibition (ER) blocked it completely, as expected (**Figure 3A**), suggesting TNF does not feedback to EGFR early after kidney injury *in vivo*. Second, we examined the effect of TNF inhibition on macrophage accumulation, which we showed earlier is significantly regulated by activation of the EGFR pathway (shown in Figure 2). Both bulk mRNA sequencing (**Figure 3B**) and mass cytometric analysis (**Figure 3C**) showed that etanercept (ET), unlike EGFR inhibition (compare to Figure 2B+C), had no effect on macrophage accumulation in the injured kidney early after AKI (Day 2). Thus, TNF inhibition does not act upstream of EGFR, nor does it drive or interfere with early EGFR-driven kidney macrophage accumulation.

**Figure 3:**
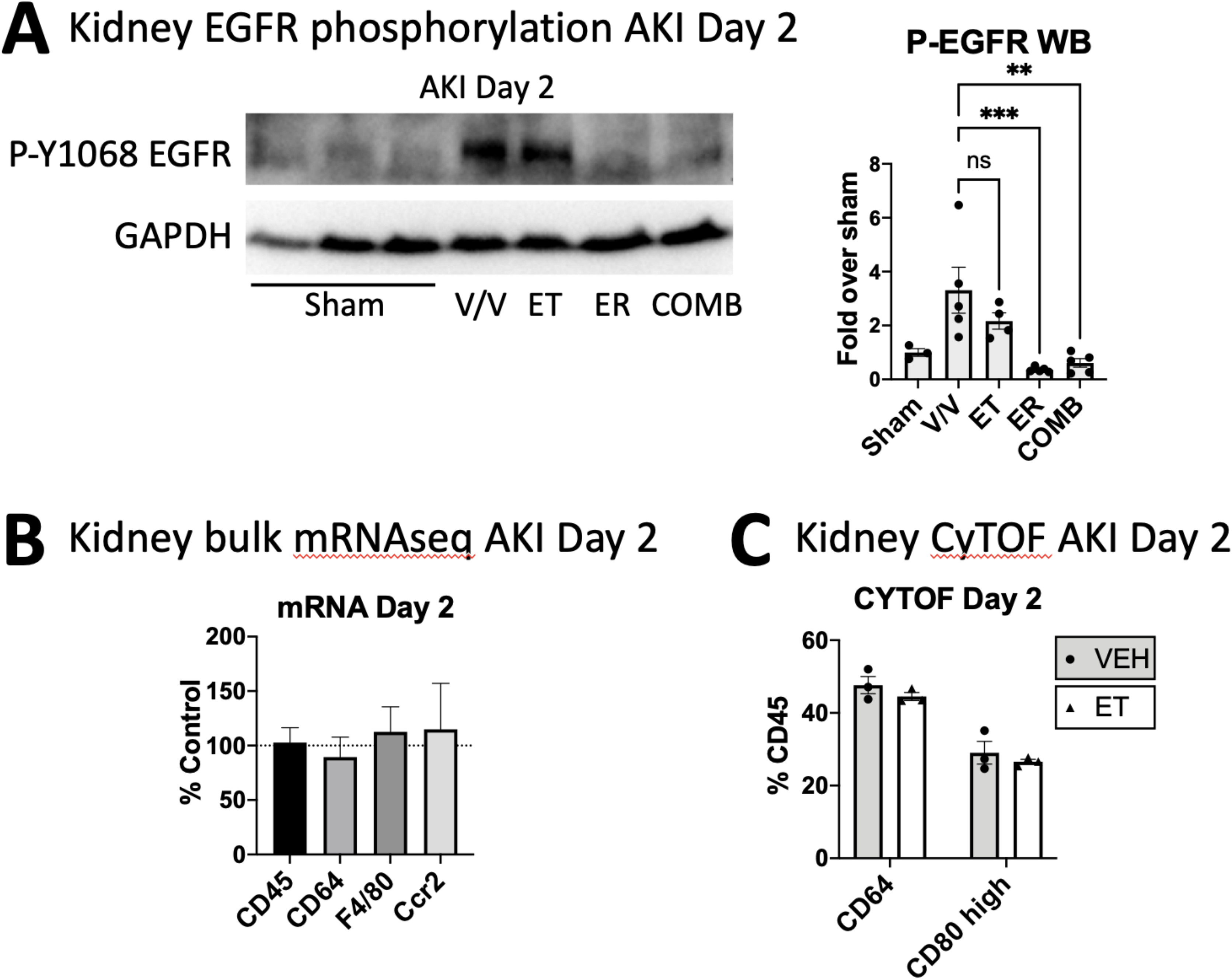
TNF inhibition does not interfere with kidney EGFR activation or macrophage accumulation in the kidney early after injury. **A**. Western blot of cortical PTC-enriched kidney lysates using anti-phospho-EGFR (Y1068) antibody. ER and COMB block PTC-EGFR phosphorylation as expected, but ET does not affect PTC-EGFR phosphorylation after AKI (n=4). **B**. Bulk mRNA sequencing analysis of kidney at Day 2 after AKI in etanercept treated AKI animals, shown as percent expression over vehicle (% control). Expression of the general immune cell marker CD45, of the macrophage marker CD64, of the macrophage/dendritic cell marker F4/80, and of immune cell recruitment receptor CCR2, are all not affected by etanercept, suggesting TNF does not significantly drive recruitment and accumulation of macrophages in the injured kidney (n=3). **C**. Mass cytometry analysis of kidney Day 2 after AKI in vehicle (VEH) or etanercept (ET) treated mice. ET does not affect the number of CD64 positive or CD80 high macrophages after AKI (n=3). **P < 0.01; ***P < 0.001; ns: non-significant

### TNF and EGFR inhibition mechanistically overlap by either downregulating identical cytokines or reducing functionally similar cytokines after AKI

As a third approach to determine whether TNF might mechanistically be connected to the profibrotic actions of EGFR signaling, we tested whether EGFR and TNF have mechanistically overlapping effects on kidney proinflammatory and profibrotic cytokine expression after AKI. We thus extended our initial proteomic cytokine analysis and examined the effect of etanercept (ET) or of the combination of erlotinib and etanercept (COMB), on cytokine expression profiles at Day 2 after AKI, similar to our analysis of mice treated with erlotinib alone (Figure 1, Supplemental Table 1+2). TNF-inhibition (ET) indeed partially overlapped in suppression of cytokines with EGFR-inhibition (ER) or combination (COMB) treatment (**Figure 4A**). Gene-ontology (GO) term analysis of all affected cytokines revealed an even more significant overlap between all treatments on the functional level (**Figure 4B**). Notably, treatments overlapped most strongly in downregulating immune response, tissue remodeling and inflammatory cell signaling events. However, and consistent with the lack of effect of TNF inhibition on kidney macrophage recruitment after injury, TNF inhibition did have significantly less negative effects on leukocyte migration and cell-cell adhesion as compared to EGFR inhibition or combination treatment (**Figure 4C**). A complete list of affected cytokines and their functional GO terms stratified by treatment condition is shown in **Supplemental Table 3**. Six cytokines were strongly downregulated by all three treatments (**Figure 4D**), suggesting they might be particularly central to shared beneficial effects between treatments in dampening the profibrotic immune response after kidney injury: TNF and interferon gamma (IFNG), an early response cytokine that orchestrates activation of the innate immune system, were the two top reduced cytokines by all treatments. Complement-factor-5 (C5), a strong chemoattractant for immune cells, including neutrophils and monocytes, p-Selectin, a mediator of the adhesion of immune cells to endothelial cells, and the innate immune system activator and regulator of cell migration Serpin E1 (also called plasminogen activator inhibitor 1) represented other key downregulated cytokines. This suggests that the severity of Type 1 inflammatory response and of innate immune cell activation, as well as the commensurate degree of neutrophil and immune cell attraction into the kidney, play important roles in AKI-to-CKD-transition. Interleukin 10 (IL10), a key antiinflammatory cytokine produced by activated immune cells was moderately reduced by all treatments, likely as a result of dampened inflammation overall, one key stimulus for IL-10 secretion. GO term analysis of the six commonly downregulated cytokines is shown in **Figure 4E**) and a complete list of functional GO terms for common downregulated cytokines is available in **Supplemental Table 4**. The similarity of the effects of EGFR and TNF inhibition on kidney cytokine expression early after AKI predict that TNF inhibition should have similar anti-fibrotic effects as did EGFR inhibition.

**Figure 4:**
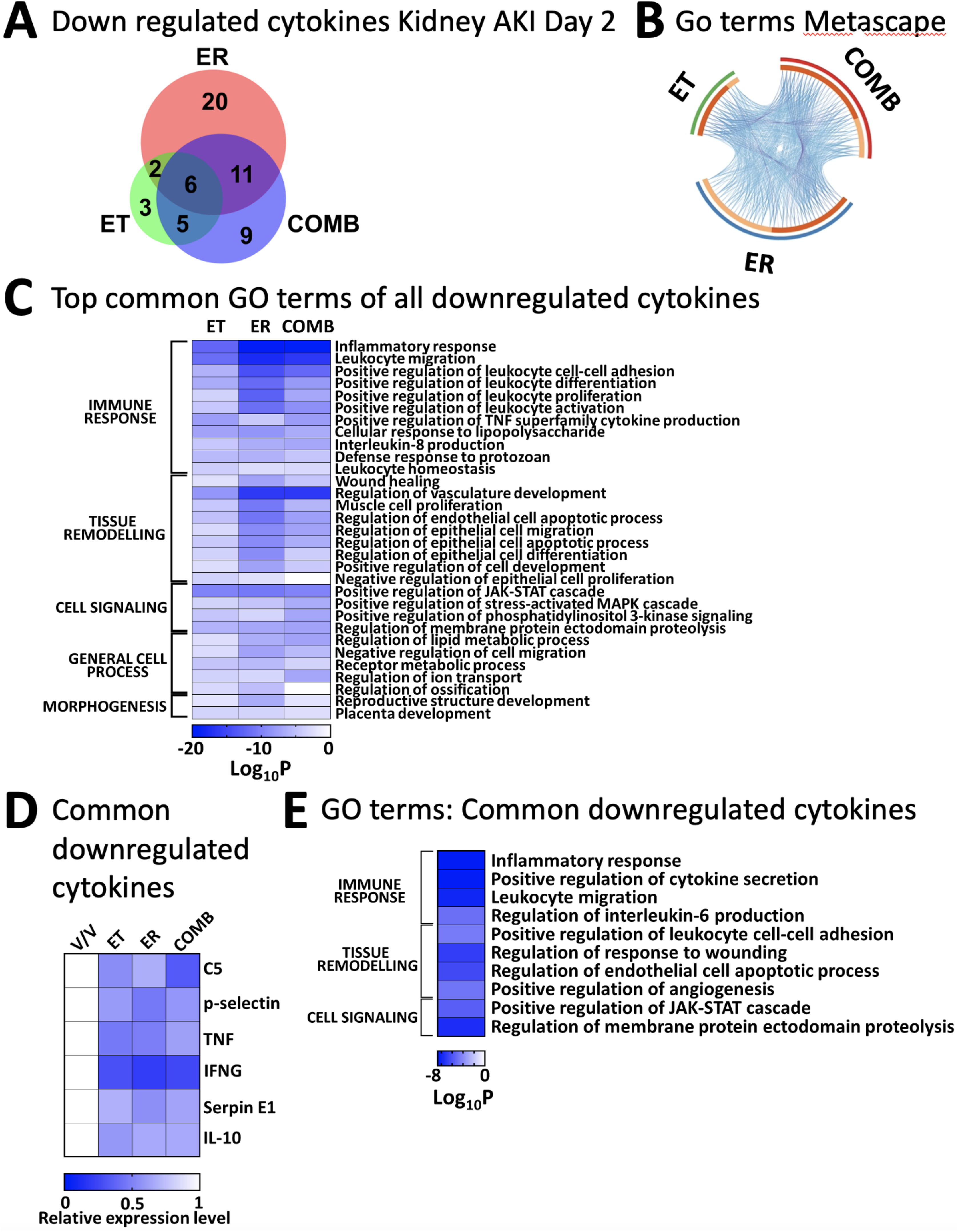
TNF and EGFR inhibition mechanistically overlap by either downregulating identical cytokines or reducing functionally similar cytokines after AKI. **A**. Venn diagram of cytokines downregulated by each treatment (erlotinib: ET; etanercept: ET; combination of etanercept and erlotinib: COMB) as compared to vehicle treated AKI animals (n=3). **B**. GO-term analysis of all downregulated cytokines. Outer circle denotes treatments; inner circle represents the downregulated cytokines: those which are common between treatments are shown in dark orange and those unique to a particular treatment are shown in light orange color. Purple curves connect identical cytokines and blue curves connect cytokines that belong to the same enriched GO term. Functional overlap between treatments is more significant than protein analysis suggests. **C**. Top common GO terms of all downregulated cytokines. Negative effects on inflammation and leukocyte migration represent top GO terms for ER and COMB and to a lesser degree for ET. **D**. Downregulated cytokines common to all treatments. TNF and IFNG, which drive type 1 inflammation and activation of the innate immune system represent top hits negatively affected by all treatments. **E**. Top common GO terms of common downregulated cytokines. Negative effects on inflammation and leukocyte migration represent top GO terms.

**Figure 5:**
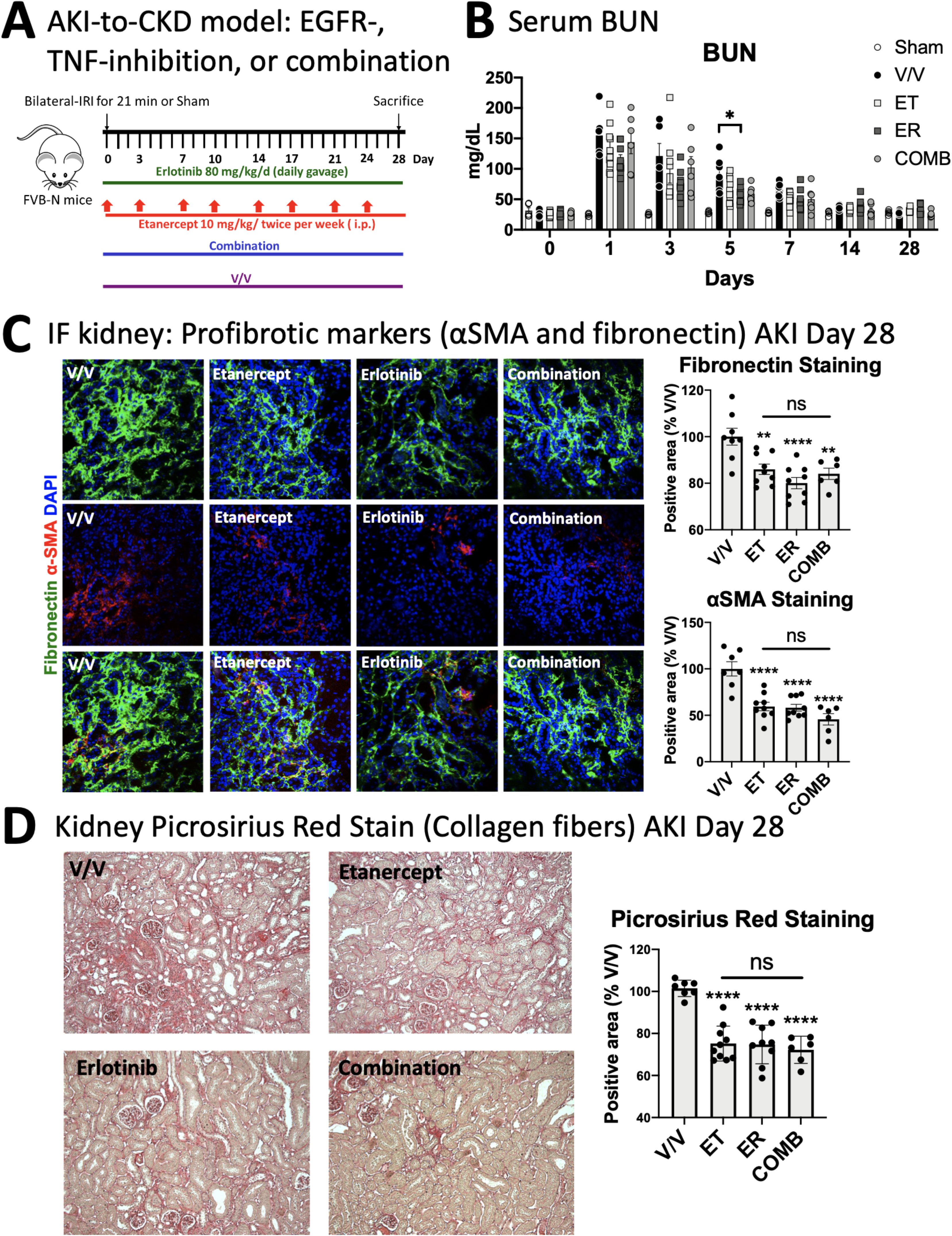
Inhibition of TNF by etanercept reduces AKI-to-CKD transition and kidney fibrosis after AKI equally effective as EGFR inhibition. **A**. Experimental scheme of mice subjected to AKI and treated with vehicle (V/V), EGFR (ER, erlotinib) or TNF-inhibition (ET, etanercept) or their combination (COMB) after AKI. **B**. Serum BUN values. BUN reaches baseline independent of treatment by Day 14 after AKI. **C**. Immunofluorescence staining of cortical area at day 28 after AKI. Upregulation of fibrosis markers Fibronectin (green) and αSMA (red) is significantly and equally reduced by all treatments as compared to vehicle treated cells. **D**. Picrosirius red staining of cortical area at day 28 after AKI. Fibrosis is significantly and equally reduced by all treatments as compared to vehicle treated cells. n = 6-10 mice/group. *P < 0.05; **P < 0.01; ****P < 0.0001; ns: non-significant.

### Inhibition of TNF by etanercept reduces AKI-to-CKD transition and kidney fibrosis after AKI equally effective as EGFR inhibition

To test whether TNF indeed mediates development of fibrosis after AKI, and to compare it to EGFR inhibition, we treated mice with TNF inhibition (ET), EGFR inhibition (ER) or their combination (COMB) for one month after AKI (**Figure 5A**). Etanercept (ET), similar to erlotinib (ER), did not affect initial injury of the kidney, or its recovery, as determined by serum BUN (**Figure 5B**) or creatinine (**Supplemental Figure 2C-Day 1 after AKI**) elevations. However, we found that, TNF-inhibition (ET) or EGFR-inhibition (ER) prevented AKI-induced fibrosis equally well, with combination treatment (COMB) providing no additional benefit, as determined by the expression of pro-fibrotic markers α-smooth-muscle-actin α-SMA and fibronectin (**Figure 5C**), as well as by picrosirius red staining of collagen fibers (**Figure 5D**). As described above for the AKI-to-CKD model (Figure 1B), BUN, creatinine or GFR have all returned to normal levels by day 28 for all the groups and are not sensitive enough to demonstrate the observed differences in fibrosis between the Vehicle and treatment groups (**Figure 5B, Supplemental Figure 2C-Day 28 after AKI, and Supplemental Figure 2D**).

Taken together, our analysis provides mechanistic insights into the anti-fibrotic effects of TNF or EGFR-inhibition in AKI-to-CKD transition. Our results reveal TNF as a key downstream effector of profibrotic EGFR activity and a key cytokine in the development of profibrotic inflammation after AKI. TNF and EGFR inhibition overlap strongly mechanistically in their ability to prevent fibrosis by (1) shared inhibition of kidney expression of several key inflammatory cytokines, including of TNF, and by (2) (largely) negatively acting on the expression of various cytokines not shared between treatments, which however share the same or similar gene functions. Thus, activation of the TNF pathway in the kidney appears to mimic many of the key pro-inflammatory and pro-fibrotic functions of EGFR pathway activation in the kidney.

## Discussion

We show that EGFR- or TNF-inhibition prevent chronic profibrotic inflammation in AKI-to-CKD transition equally effectively, because TNF acts as a major effector downstream of profibrotic EGFR signaling after AKI. EGFR-inhibition reduces recruitment of profibrotic TNF-producing immune cells early after AKI, limiting profibrotic inflammation, whereas TNF-inhibition neutralizes soluble TNF produced by these profibrotic immune cells and possibly other immune or kidney resident cells, and blunts kidney inflammation and the production of cytokines. But this does not entirely explain the proinflammatory and profibrotic action of both pathways. While there is significant overlap between cytokines activated by the EGFR and TNF pathway, including TNF and INFG, both pathways also regulate separate, yet functionally overlapping, sets of cytokines.

Consistent with our data, TNF-inhibition has also been shown to decrease glomerular inflammation and glomerulosclerosis (glomerular fibrosis) in diabetic nephropathy. In diabetic Akita mice, anti-TNF-antibody reduced glomerular macrophages, glomerulosclerosis, albuminuria and prevented reduction in glomerular filtration rate ^32^, consistent with an anti-fibrotic effect of TNF-inhibition in inflamed glomeruli. This study also highlighted the importance of TNF expression in macrophages for induction of fibrosis in the injured kidney, as suggested by our findings. In the streptozosin (STZ) diabetic model, TNF-macrophage-KO reduced BUN and creatinine elevations, glomerular macrophage accumulation, albuminuria and TNF mRNA/protein in the kidney ^32^. The AKI kidney snRNAseq data we used to assess cellular TNF sources after AKI ^26^ did not include dendritic cells, and published data suggests that dendritic cells in the kidney could also represent possible sources of profibrotic TNF after AKI. Dong et al. showed that dendritic cells represent a major source of TNF early after renal IRI (24hours) ^33^. However, the role of dendritic cells in kidney fibrosis has not been studied to our knowledge. Yet, dendritic cells and dendritic cell-derived TNF have been associated with fibrosis in the liver ^34^ and the lung ^35,36^.

Complicating any analysis of TNF targeting agents is the fact that they not only bind to soluble TNF, which preferentially activates TNFR1, a receptor that mediates the classic proinflammatory injurious actions of soluble TNF. In addition, some TNF targeting agents, including etanercept, can bind to uncleaved transmembrane pro-TNF, which strongly activates TNFR2. TNFR2 is expressed in immune cells and requires cell-cell contact for activation, with the ligand and receptor expressed in opposing cells ^37^. TNFR2 activation by pro-TNF can have either proinflammatory or anti-inflammatory/pro-repair effects, depending on the cellular context ^38^. pro-TNF:TNFR2 interaction could be disturbed when TNF-targeting agents bind to transmembrane pro-TNF, possibly explaining some of our results using etanercept. Results in TNF-global-knockout mice also have to be interpreted carefully, because they are missing soluble and pro-TNF. This possibly explains why some of the earlier studies of TNF-inhibition ^17^ or of TNF-global-knockout mice ^39^ yielded divergent results from our studies. In contrast to the study using UUO and pegylated-TNFR1 in rats ^17^, etanercept did not reduce early kidney injury in our studies.

We previously showed that soluble TNF induces moderate ADAM17-mediated release of AREG in PTC *in vitro* ^4^, allowing the possibility that TNF might act upstream of profibrotic PTC-EGFR signaling *in vivo*. However, TNF-inhibition had no effect on kidney EGFR activation in our current study, suggesting that TNF-induced EGFR activation does not contribute to the development of AKI-induced fibrosis *in vivo*.

Although AKI-to-CKD transition with progressive loss of GFR is by now a well-established clinical concept ^1,2,40–42^, its dynamic development has to our knowledge not been shown on the GFR level in a mouse model. Others confirmed our findings on early injury resolution with normalization of biomarkers over 14 days in the bilateral renal IRI model, but found that Creatinine was still normal at 6 months and only elevated by 12 months after AKI ^43^, revealing higher sensitivity of GFR measurements in our studies, which detected GFR decreases 3-4 months after AKI, which were not revealed by serum BUN or creatinine measurements.

Our studies raise the possibility that TNF or EGFR-inhibition could be used clinically in kidney disease. EGFR-inhibition has largely been used in the treatment of cancer. Because EGFR signaling is important for homeostasis, proliferation and wound healing in many epithelial cells, EGFR-inhibition typically leads to diarrhea, wound healing defects and skin infections. Blunting epithelial repair by EGFR-inhibition leading to delayed renal recovery after AKI has been observed in mice ^24^. For these reasons, long-term EGFR-inhibition would likely not represent a viable therapeutic strategy clinically. TNF-inhibition on the other hand has been used in various autoimmune diseases for long periods of time with significant benefits that are weighed against potentially serious adverse effects and risk of morbidity/mortality due to disease progression. Typical adverse effects include injection site reactions, infusion reactions (both usually manageable), neutropenia (usually mild), skin lesions and infections; autoimmune diseases develop rarely ^44–46^. The risk of heart failure remains unclear, as combined analysis of the two performed relevant trials showed no effect on death or new heart failure, with only few patients possibly experiencing heart failure exacerbations ^47^. Several described that can develop in response to TNF inhibitors are rather rare, including reports of such diseases affecting the kidney itself. Thus, in our view, the opportunity for TNF-inhibition or EGFR-inhibition in kidney patients could center on short- or medium-term treatment used relatively early after onset of AKI when AKI-to-CKD transition could be prevented or blunted. TNF-inhibition would likely be preferrable to EGFR-inhibition based on potential adverse effects. Given the high yearly mortality of CKD patients this could be a justifiable approach and be performed at reasonable risk.

## Acknowledgements

We thank Pfizer for the provision of murine etanercept. Dr. Herrlich was supported by National Institute of Diabetes and Digestive and Kidney Diseases (R01DK108947 and R01DK121200) and Dr. Kefaloyianni was supported by the American Heart Association (Career Development Award 20CDA35320006) and the American Society of Nephrology (KidneyCure Carl W. Gottschalk Research Scholar Grant).

## Supplemental Data

### Table of Contents

**Supplemental Figure 1:** AKI-to-CKD model: Assessment of traditional markers of kidney function, BUN (**A**) and creatinine (**B**), over 6 months.

**Supplemental Figure 2:** Injected fluorescently-labelled murine etanercept is detectable in (**A**) kidney interstitium of sham and AKI mice, and (**B**) partially overlaps with CD31 expression, which marks endothelial cells, showing that some of the labelled murine etanercept is clearly located outside of vessels. (**C**) Serum creatinine values at Day 1 and Day 28 are equal in all treatment groups. (**D**) GFR measurements at Day 28 are equal in all treatment groups.

**Supplemental Table 1:** GO term analysis for all downregulated cytokines by EGFR inhibition with erlotinib (ER).

**Supplemental Table 2:** Summary of enrichment analysis of transcription factors known to regulate cytokines downregulated by EGFR inhibition with erlotinib (ER).

**Supplemental Table 3:** GO term analysis for all downregulated cytokines stratified by treatment condition.

**Supplemental Table 4:** GO term analysis for downregulated cytokines common between all treatment conditions.

