## Supplemental Files for "EGFR-driven profibrotic signals in AKI-to-CKD transition are blocked by TNF inhibition: Opportunities for etanercept treatment"

**Supplemental Figure 1:** AKI-to-CKD model: Assessment of traditional markers of kidney function, BUN (**A**) and creatinine (**B**), over 6 months.

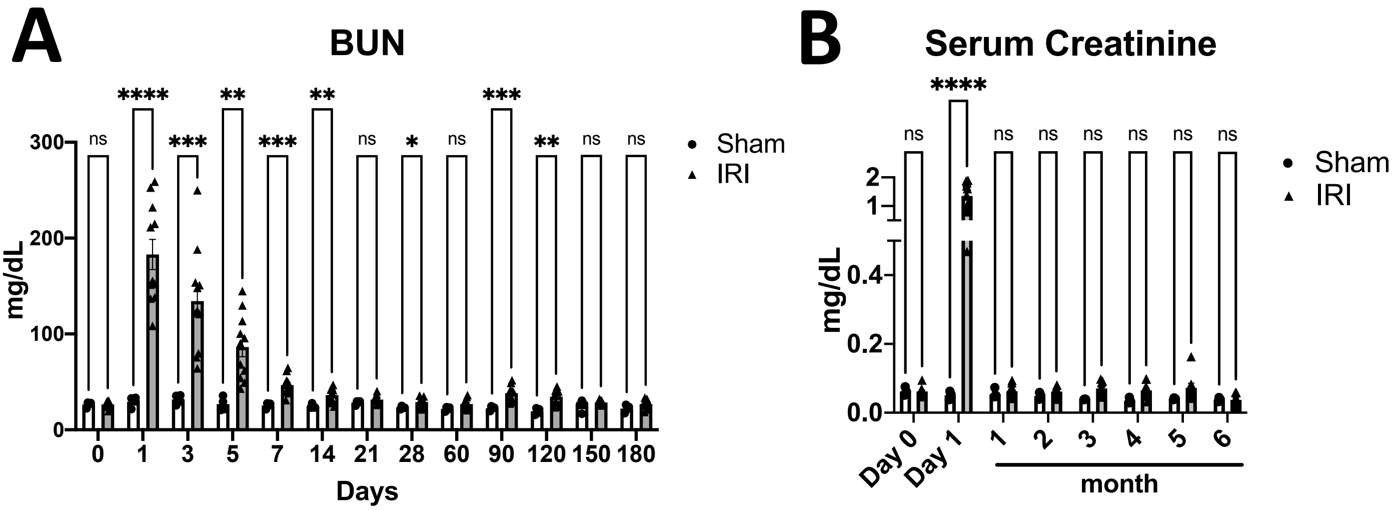

**Supplemental Figure 2:** Injected fluorescently-labelled murine etanercept is detectable in (**A**) kidney interstitium of sham and AKI mice, and (**B**) partially overlaps with CD31 expression, which marks endothelial cells, showing that some of the labelled murine etanercept is clearly located outside of vessels. (**C**) Serum creatinine values at Day 1 and Day 28 are equal in all treatment groups. (**D**) GFR measurements at Day 28 are equal in all treatment groups.

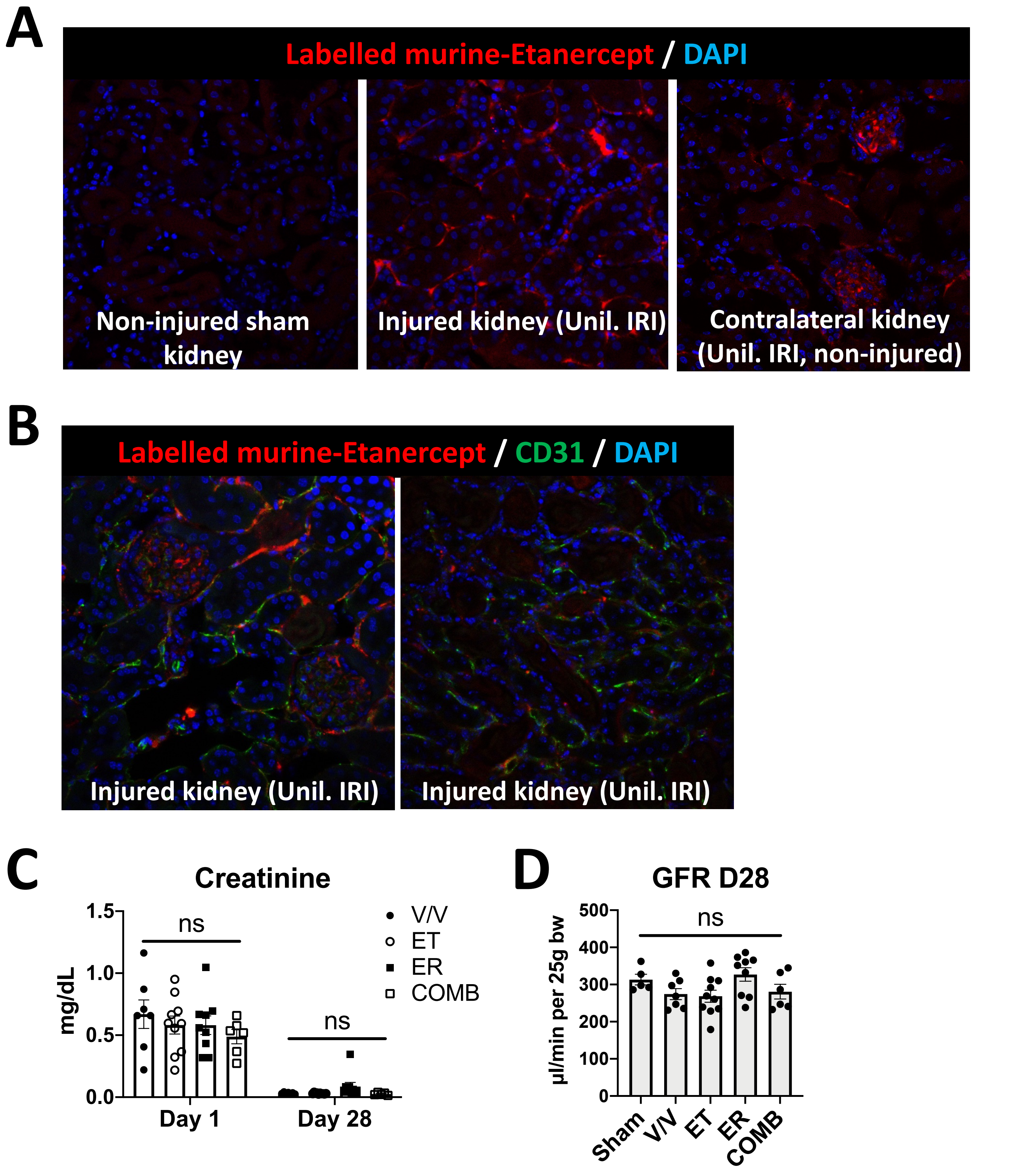

**Supplemental Table 1:** GO term analysis for all downregulated cytokines by EGFR inhibition with erlotinib (ER).

| Category | Term | Description | _LogP_ER | Gene count | Genes |
|---|---|---|---|---|---|
| GO BP | GO:0006954 | inflammatory response | -19.9094 | 30 | Adipoq\|Ahsg\|Crp\|Csf1\|Cxcl1\|Hc\|Hgf\|Ifng\|Il10\|Il12b\|Il1a\|Il1b\|Il1rn\|Il4\|Ldlr\|Lep\|Cxcl9\|Serpine1\|Ccl21a\|Ccl2\|Ccl22\|Ccl3\|Ccl4\|Cxcl5\|Serpinf1\|Selp\|Tnf\|Ccn4\|Ccl19\|Il23a |
| GO BP | GO:0050900 | leukocyte migration | -17.8334 | 24 | Csf1\|Gas6\|Cxcl1\|Hc\|Ifng\|Il1a\|Il1b\|Il4\|Lep\|Cxcl9\|Serpine1\|Ccl21a\|Ccl2\|Ccl22\|Ccl3\|Ccl4\|Cxcl5\|Selp\|Tnf\|Vcam1\|Vegfa\|Ccl19\|Cxcl16\|Il23a |
| GO BP | GO:0019221 | cytokine-mediated signaling pathway | -13.047 | 22 | Adipoq\|Angpt1\|Csf1\|Gas6\|Cxcl1\|Ifng\|Il12b\|Il15\|Il1a\|Il1b\|Il1rn\|Lep\|Cxcl9\|Ccl21a\|Ccl2\|Ccl22\|Ccl3\|Ccl4\|Cxcl5\|Thpo\|Tnf\|Ccl19 |
| GO BP | GO:1903708 | positive regulation of hemopoiesis | -12.589 | 18 | Csf1\|Csf3\|Dlk1\|Gas6\|Cxcl1\|Ifng\|Il12b\|Il15\|Il1a\|Il1b\|Il4\|Lep\|Lif\|Ccl3\|Thpo\|Tnf\|Ccl19\|Il23a |
| GO BP | GO:1901342 | regulation of vasculature development | -16.3137 | 19 | Angpt2\|Fgf1\|Hc\|Hgf\|Ifng\|Il10\|Il12b\|Il1a\|Il1b\|Lep\|Lif\|Serpine1\|Prl2c2\|Ccl2\|Serpinf1\|Tnf\|Vegfa\|Angptl3\|Cd160 |
| GO BP | GO:0050731 | positive regulation of peptidyl-tyrosine phosphorylation | -13.2519 | 15 | Adipoq\|Angpt1\|Csf3\|Gas6\|Hgf\|Ifng\|Il11\|Il12b\|Il15\|Il4\|Lep\|Lif\|Tnf\|Vegfa\|Il23a |
| GO BP | GO:0033002 | muscle cell proliferation | -10.8069 | 12 | Adipoq\|Angpt1\|Fgf1\|Ifng\|Igfbp5\|Il10\|Il12b\|Il15\|Rbp4\|Tnf\|Vegfa\|Ccn4 |
| GO BP | GO:2000351 | regulation of endothelial cell apoptotic process | -11.0221 | 7 | Angpt1\|Gas6\|Il10\|Il4\|Serpine1\|Tnf\|Cd160 |
| GO BP | GO:0071222 | cellular response to lipopolysaccharide | -8.60998 | 12 | Cxcl1\|Ifng\|Il10\|Il12b\|Il1a\|Il1b\|Cxcl9\|Serpine1\|Ccl2\|Cxcl5\|Tnf\|Cxcl16 |
| GO BP | GO:0006959 | humoral immune response | -6.10929 | 13 | Cfd\|Crp\|Cxcl1\|Hc\|Ifng\|Il1b\|Cxcl9\|Apcs\|Ccl2\|Ccl22\|Cxcl5\|Tnf\|Ccl19 |
| GO BP | GO:0001659 | temperature homeostasis | -8.82196 | 10 | Adipoq\|Il15\|Il1a\|Il1b\|Il1rn\|Il4\|Lcn2\|Lep\|Tnf\|Vegfa |
| GO BP | GO:0006953 | acute-phase response | -9.87474 | 6 | Ahsg\|Crp\|Il1a\|Il1b\|Il1rn\|Tnf |
| GO BP | GO:0007169 | transmembrane receptor protein tyrosine kinase signaling pathway | -6.99063 | 14 | Adipoq\|Angpt1\|Angpt2\|Ahsg\|Csf1\|Fgf1\|Hgf\|Igfbp2\|Igfbp5\|Il12b\|Il1b\|Lep\|Ccl2\|Vegfa |
| GO BP | GO:0010632 | regulation of epithelial cell migration | -9.34714 | 9 | Angpt1\|Fgf1\|Ifng\|Il4\|Serpine1\|Prl2c2\|Serpinf1\|Tnf\|Vegfa |
| GO BP | GO:0045834 | positive regulation of lipid metabolic process | -9.09926 | 9 | Adipoq\|Fgf1\|Ifng\|Il1a\|Il1b\|Ldlr\|Tnf\|Ccl19\|Angptl3 |
| GO BP | GO:0034113 | heterotypic cell-cell adhesion | -9.12331 | 6 | Adipoq\|Il10\|Il1b\|Il1rn\|Tnf\|Vcam1 |
| GO BP | GO:0030225 | macrophage differentiation | -7.87213 | 6 | Adipoq\|Csf1\|Ifng\|Il15\|Lif\|Vegfa |
| GO BP | GO:0010720 | positive regulation of cell development | -7.48878 | 14 | Adipoq\|Crp\|Csf1\|Dkk1\|Hgf\|Ifng\|Il1b\|Lif\|Prl2c2\|Cxcl5\|Serpinf1\|Tnf\|Vegfa\|Ccn4 |
| GO BP | GO:0048771 | tissue remodeling | -5.71951 | 9 | Angpt2\|Ahsg\|Dlk1\|Igfbp5\|Il1a\|Lep\|Lif\|Vegfa\|Il23a |
| GO BP | GO:0014065 | phosphatidylinositol 3-kinase signaling | -3.98861 | 6 | Csf3\|Hgf\|Lep\|Selp\|Tnf\|Cd160 |
| GO BP | GO:0042060 | wound healing | -7.401 | 9 | Crp\|Fgf1\|Gas6\|Il1a\|Serpine1\|Selp\|Tnf\|Vegfa\|Ccn4 |
| GO BP | GO:0043086 | negative regulation of catalytic activity | -7.29026 | 11 | Adipoq\|Ahsg\|Gas6\|Ifng\|Il1b\|Serpine1\|Apcs\|Serpinf1\|Tnf\|Vegfa\|Angptl3 |
| GO BP | GO:0048608 | reproductive structure development | -6.67768 | 11 | Angpt1\|Ahsg\|Il10\|Il1a\|Lep\|Lif\|Serpine1\|Rbp4\|Serpinf1\|Vcam1\|Vegfa |
| GO BP | GO:0043269 | regulation of ion transport | -3.3508 | 10 | Cxcl1\|Ifng\|Il1b\|Il1rn\|Il4\|Lep\|Serpine1\|Ccl2\|Ccl4\|Tnf |
| GO BP | GO:0030278 | regulation of ossification | -5.23991 | 8 | Ahsg\|Csf1\|Dkk1\|Dlk1\|Igfbp5\|Tnf\|Vegfa\|Ccn4 |
| GO BP | GO:0042832 | defense response to protozoan | -6.20523 | 4 | Ifng\|Il10\|Il12b\|Il4 |
| GO BP | GO:0010876 | lipid localization | -5.04559 | 9 | Adipoq\|Crp\|Il1a\|Il1b\|Ldlr\|Lep\|Rbp4\|Tnf\|Angptl3 |
| GO BP | GO:0031294 | lymphocyte costimulation | -4.56307 | 4 | Il4\|Ccl19\|Tnfsf13b\|Cd160 |
| GO BP | GO:0001959 | regulation of cytokine-mediated signaling pathway | -5.72004 | 5 | Adipoq\|Angpt1\|Csf1\|Gas6\|Il1rn |
| GO BP | GO:0061061 | muscle structure development | -5.28509 | 11 | Angpt1\|Dkk1\|Igfbp5\|Il4\|Lif\|Cxcl9\|Prl2c2\|Rbp4\|Tnf\|Vegfa\|Ccn4 |
| GO BP | GO:0010743 | regulation of macrophage derived foam cell differentiation | -4.96302 | 3 | Adipoq\|Crp\|Csf1 |
| GO BP | GO:2000648 | positive regulation of stem cell proliferation | -3.76434 | 4 | Cxcl1\|Prl2c2\|Thpo\|Vegfa |
| GO BP | GO:0001890 | placenta development | -3.49003 | 5 | Il10\|Lep\|Lif\|Serpine1\|Vcam1 |
| GO BP | GO:0045661 | regulation of myoblast differentiation | -3.61484 | 3 | Cxcl9\|Prl2c2\|Tnf |

**Supplemental Table 2:** Summary of enrichment analysis of transcription factors known to regulate cytokines downregulated by EGFR inhibition with erlotinib (ER).

| Category | Term | Description | _LogP_ER | Gene count | Genes |
|---|---|---|---|---|---|
| TRRUST | TRR01349 | Regulated by: Sp3 | -2.9 | 3 | Angpt2\|Hgf\|Il10 |
| TRRUST | TRR01347 | Regulated by: Sp1 | -3.2 | 5 | Hgf\|Il10\|Serpine1\|Ccl2\|Tnf |
| TRRUST | TRR01462 | Regulated by: Tcf3 | -3.7 | 3 | Adipoq\|Ccl21a\|Vcam1 |
| TRRUST | TRR01316 | Regulated by: Smad4 | -4.1 | 3 | Il10\|Vegfa\|Tnfsf13b |
| TRRUST | TRR00293 | Regulated by: Egr1 | -4.2 | 5 | Il1b\|Il4\|Serpine1\|Ccl2\|Tnf |
| TRRUST | TRR00090 | Regulated by: Bcl3 | -4.7 | 3 | Ifng\|Tnf\|Il23a |
| TRRUST | TRR00437 | Regulated by: Gata3 | -5.2 | 3 | Ifng\|Il10\|Il4 |
| TRRUST | TRR01535 | Regulated by: Wt1 | -5.2 | 3 | Csf1\|Il10\|Vegfa |
| TRRUST | TRR01315 | Regulated by: Smad3 | -5.3 | 4 | Serpine1\|Vegfa\|Tnfsf13b\|Il23a |
| TRRUST | TRR00638 | Regulated by: Irf8 | -6.3 | 4 | Ifng\|Il12b\|Il1b\|Tnf |
| TRRUST | TRR00631 | Regulated by: Irf1 | -6.3 | 4 | Ifng\|Il12b\|Il4\|Tnf |
| TRRUST | TRR01368 | Regulated by: Stat3 | -7.5 | 6 | Igfbp5\|Il10\|Lif\|Tnf\|Vegfa\|Il23a |
| TRRUST | TRR00939 | Regulated by: Nfatc2 | -8.3 | 5 | Ifng\|Il10\|Il12b\|Il4\|Tnf |
| TRRUST | TRR00634 | Regulated by: Irf4 | -8.4 | 4 | Il10\|Il1b\|Il4\|Cxcl9 |
| TRRUST | TRR01126 | Regulated by: Pparg | -8.4 | 6 | Adipoq\|Csf1\|Fgf1\|Ifng\|Il4\|Serpine1 |
| TRRUST | TRR00652 | Regulated by: Jun | -8.9 | 8 | Angpt2\|Cxcl1\|Ifng\|Il1b\|Il4\|Serpine1\|Ccl2\|Tnf |
| TRRUST | TRR01189 | Regulated by: Rel | -9.5 | 6 | Ifng\|Il12b\|Il1b\|Il4\|Tnf\|Il23a |
| TRRUST | TRR01190 | Regulated by: Rela | -13 | 10 | Csf1\|Ifng\|Il10\|Il12b\|Il4\|Selp\|Tnf\|Vcam1\|Tnfsf13b\|Il23a |
| TRRUST | TRR00951 | Regulated by: Nfkb1 | -20 | 16 | Adipoq\|Csf1\|Ifng\|Il10\|Il12b\|Il1a\|Il1b\|Il4\|Lcn2\|Lif\|Selp\|Tnf\|Vcam1\|Vegfa\|Tnfsf13b\|Il23a |

**Supplemental Table 3:** GO term analysis for all downregulated cytokines stratified by treatment condition.

| Category | Term | Description | _LogP_ET | _LogP_ER | _LogP_COMB | Log(q) | Gene count | Genes |
|---|---|---|---|---|---|---|---|---|
| GO BP | GO:0006954 | Inflammatory response | -12.344 | -19.9094 | -19.556 | -23.92696374 | 30 | Adipoq\|Ahsg\|Crp\|Csf1\|Cxcl1\|Hc\|Hgf\|Ifng\|Il10\|Il12b\|Il1a\|Il1b\|Il1rn\|Il4\|Ldlr\|Lep\|Cxcl9\|Serpine1\|Ccl21a\|Ccl2\|Ccl22\|Ccl3\|Ccl4\|Cxcl5\|Serpinf1\|Selp\|Tnf\|Ccn4\|Ccl19\|Il23a |
| GO BP | GO:0050900 | Leukocyte migration | -11.8418 | -17.8334 | -16.3962 | -23.18010055 | 24 | Csf1\|Gas6\|Cxcl1\|Hc\|Ifng\|Il1a\|Il1b\|Il4\|Lep\|Cxcl9\|Serpine1\|Ccl21a\|Ccl2\|Ccl22\|Ccl3\|Ccl4\|Cxcl5\|Selp\|Tnf\|Vcam1\|Vegfa\|Ccl19\|Cxcl16\|Il23a |
| GO BP | GO:1903039 | Positive regulation of leukocyte cell-cell adhesion | -6.26421 | -14.272 | -12.2133 | -15.48177512 | 16 | Ifng\|Igfbp2\|Il12b\|Il15\|Il1a\|Il1b\|Il4\|Lep\|Ccl2\|Selp\|Tnf\|Vcam1\|Ccl19\|Tnfsf13b\|Cd160\|Il23a |
| GO BP | GO:1902107 | Positive regulation of leukocyte differentiation | -6.70637 | -11.9784 | -8.11191 | -13.78842401 | 14 | Csf1\|Dlk1\|Gas6\|Ifng\|Il12b\|Il15\|Il1a\|Il1b\|Il4\|Lif\|Ccl3\|Tnf\|Ccl19\|Il23a |
| GO BP | GO:0070665 | Positive regulation of leukocyte proliferation | -3.6845 | -12.7178 | -7.02909 | -13.24880026 | 13 | Csf1\|Ifng\|Igfbp2\|Il12b\|Il15\|Il1a\|Il1b\|Il4\|Lep\|Vcam1\|Ccl19\|Tnfsf13b\|Il23a |
| GO BP | GO:0002696 | Positive regulation of leukocyte activation | -4.62429 | -11.6246 | -8.91063 | -11.90951525 | 17 | Gas6\|Ifng\|Igfbp2\|Il10\|Il12b\|Il15\|Il1a\|Il1b\|Il4\|Lep\|Ccl2\|Tnf\|Vcam1\|Ccl19\|Tnfsf13b\|Cd160\|Il23a |
| GO BP | GO:1903557 | Positive regulation of tumor necrosis factor superfamily cytokine production | -7.87983 | -4.33674 | -7.99144 | -8.852529374 | 9 | Ifng\|Il12b\|Il1a\|Lep\|Ccl2\|Ccl3\|Ccl4\|Ccl19\|Il23a |
| GO BP | GO:0071222 | Cellular response to lipopolysaccharide | -7.31876 | -8.60998 | -6.74276 | -8.765524224 | 12 | Cxcl1\|Ifng\|Il10\|Il12b\|Il1a\|Il1b\|Cxcl9\|Serpine1\|Ccl2\|Cxcl5\|Tnf\|Cxcl16 |
| GO BP | GO:0032637 | Interleukin-8 production | -4.59791 | -6.53019 | -7.05034 | -5.463100492 | 6 | Adipoq\|Crp\|Il1b\|Lep\|Serpine1\|Tnf |
| GO BP | GO:0042832 | Defense response to protozoan | -5.56564 | -6.20523 | -4.66894 | -4.772458709 | 4 | Ifng\|Il10\|Il12b\|Il4 |
| GO BP | GO:0001776 | Leukocyte homeostasis | -4.0431 | -2.87106 | -3.1651 | -3.331914639 | 5 | Hc\|Ifng\|Ccl2\|Cxcl5\|Tnfsf13b |
| GO BP | GO:0042060 | Wound healing | -2.4913 | -7.401 | -4.63684 | -5.765153273 | 9 | Crp\|Fgf1\|Gas6\|Il1a\|Serpine1\|Selp\|Tnf\|Vegfa\|Ccn4 |
| GO BP | GO:1901342 | Regulation of vasculature development | -8.35609 | -16.3137 | -16.464 | -16.30939662 | 19 | Angpt2\|Fgf1\|Hc\|Hgf\|Ifng\|Il10\|Il12b\|Il1a\|Il1b\|Lep\|Lif\|Serpine1\|Prl2c2\|Ccl2\|Serpinf1\|Tnf\|Vegfa\|Angptl3\|Cd160 |
| GO BP | GO:0033002 | Muscle cell proliferation | -4.55836 | -10.8069 | -5.90532 | -9.660839587 | 12 | Adipoq\|Angpt1\|Fgf1\|Ifng\|Igfbp5\|Il10\|Il12b\|Il15\|Rbp4\|Tnf\|Vegfa\|Ccn4 |
| GO BP | GO:2000351 | Regulation of endothelial cell apoptotic process | -5.00438 | -11.0221 | -7.73678 | -8.812265517 | 7 | Angpt1\|Gas6\|Il10\|Il4\|Serpine1\|Tnf\|Cd160 |
| GO BP | GO:0010632 | Regulation of epithelial cell migration | -3.13089 | -9.34714 | -7.31378 | -7.391008858 | 9 | Angpt1\|Fgf1\|Ifng\|Il4\|Serpine1\|Prl2c2\|Serpinf1\|Tnf\|Vegfa |
| GO BP | GO:1904035 | Regulation of epithelial cell apoptotic process | -4.30405 | -9.34245 | -6.55658 | -7.391008858 | 7 | Angpt1\|Gas6\|Il10\|Il4\|Serpine1\|Tnf\|Cd160 |
| GO BP | GO:0010720 | Positive regulation of cell development | -2.81308 | -7.48878 | -4.2741 | -6.67459605 | 14 | Adipoq\|Crp\|Csf1\|Dkk1\|Hgf\|Ifng\|Il1b\|Lif\|Prl2c2\|Cxcl5\|Serpinf1\|Tnf\|Vegfa\|Ccn4 |
| GO BP | GO:0050680 | Negative regulation of epithelial cell proliferation | -3.71865 | -2.55852 | 0 | -2.660636053 | 3 | Dlk1\|Il12b\|Tnf |
| GO BP | GO:0046427 | Positive regulation of JAK-STAT cascade | -9.97903 | -10.788 | -9.81382 | -12.08369482 | 11 | Ifng\|Il10\|Il12b\|Il15\|Il4\|Lep\|Lif\|Prl2c2\|Cxcl5\|Tnf\|Il23a |
| GO BP | GO:0032874 | Positive regulation of stress-activated MAPK cascade | -3.534 | -4.76456 | -6.72594 | -5.859655477 | 8 | Dkk1\|Il1a\|Il1b\|Il1rn\|Lep\|Tnf\|Vegfa\|Ccl19 |
| GO BP | GO:0014068 | Positive regulation of phosphatidylinositol 3-kinase signaling | -4.6332 | -3.44596 | -7.10977 | -5.527131844 | 6 | Angpt1\|Csf3\|Hgf\|Lep\|Selp\|Tnf |
| GO BP | GO:0051043 | Regulation of membrane protein ectodomain proteolysis | -6.16844 | -7.0205 | -7.43524 | -5.795540441 | 4 | Ifng\|Il10\|Il1b\|Tnf |
| GO BP | GO:0019216 | Regulation of lipid metabolic process | -2.58883 | -6.47654 | -7.31503 | -5.533237876 | 10 | Adipoq\|Fgf1\|Ifng\|Il1a\|Il1b\|Ldlr\|Lep\|Tnf\|Ccl19\|Angptl3 |
| GO BP | GO:0030336 | Negative regulation of cell migration | -2.91997 | -7.3639 | -5.49342 | -5.73279668 | 8 | Adipoq\|Hc\|Igfbp5\|Il1rn\|Il4\|Serpine1\|Serpinf1\|Tnf |
| GO BP | GO:0043112 | Receptor metabolic process | -4.74103 | -5.56036 | -3.55592 | -4.153143813 | 7 | Adipoq\|Angpt1\|Dkk1\|Ifng\|Il10\|Tnf\|Vegfa |
| GO BP | GO:0043269 | Regulation of ion transport | -3.71436 | -3.3508 | -7.08748 | -5.50752653 | 10 | Cxcl1\|Ifng\|Il1b\|Il1rn\|Il4\|Lep\|Serpine1\|Ccl2\|Ccl4\|Tnf |
| GO BP | GO:0030278 | Regulation of ossification | -3.09914 | -5.23991 | 0 | -4.843585494 | 8 | Ahsg\|Csf1\|Dkk1\|Dlk1\|Igfbp5\|Tnf\|Vegfa\|Ccn4 |
| GO BP | GO:0048608 | Reproductive structure development | -2.25316 | -6.67768 | -2.25787 | -5.455071122 | 11 | Angpt1\|Ahsg\|Il10\|Il1a\|Lep\|Lif\|Serpine1\|Rbp4\|Serpinf1\|Vcam1\|Vegfa |
| GO BP | GO:0001890 | Placenta development | -3.51182 | -3.49003 | -2.64774 | -2.566615937 | 5 | Il10\|Lep\|Lif\|Serpine1\|Vcam1 |

**Supplemental Table 4:** GO term analysis for downregulated cytokines common between all treatment conditions.

| Category | Term | Description | LogP | Log(q) | Gene count | Genes |
|---|---|---|---|---|---|---|
| GO BP | GO:0006954 | inflammatory response | -8.671 | -4.74101 | 6 | Hc\|Ifng\|Il10\|Serpine1\|Selp\|Tnf |
| GO BP | GO:0050715 | positive regulation of cytokine secretion | -8.63693 | -4.74101 | 4 | Hc\|Ifng\|Il10\|Tnf |
| GO BP | GO:0050900 | leukocyte migration | -8.02344 | -4.47195 | 5 | Hc\|Ifng\|Serpine1\|Selp\|Tnf |
| GO BP | GO:0032675 | regulation of interleukin-6 production | -5.06583 | -2.70139 | 3 | Ifng\|Il10\|Tnf |
| GO BP | GO:1903039 | positive regulation of leukocyte cell-cell adhesion | -4.62099 | -2.40631 | 3 | Ifng\|Selp\|Tnf |
| GO BP | GO:1903034 | regulation of response to wounding | -6.90887 | -3.75331 | 4 | Il10\|Serpine1\|Selp\|Tnf |
| GO BP | GO:2000351 | regulation of endothelial cell apoptotic process | -6.44329 | -3.49735 | 3 | Il10\|Serpine1\|Tnf |
| GO BP | GO:0045766 | positive regulation of angiogenesis | -4.83116 | -2.54802 | 3 | Hc\|Il10\|Serpine1 |
| GO BP | GO:0046427 | positive regulation of JAK-STAT cascade | -5.61685 | -3.092 | 3 | Ifng\|Il10\|Tnf |
| GO BP | GO:0051043 | regulation of membrane protein ectodomain proteolysis | -7.61243 | -4.26058 | 3 | Ifng\|Il10\|Tnf |
